# Propranolol reduces sarcoma growth and enhances the response to anti-CTLA4 therapy by modulating the tumor microenvironment

**DOI:** 10.1101/2021.03.11.434711

**Authors:** Klaire Yixin Fjæstad, Anne Mette Askehøj Rømer, Victor Goitea, Astrid Zedlitz Johansen, Marie-Louise Thorseth, Marco Carretta, Lars Henning Engelholm, Lars Grøntved, Niels Junker, Daniel Hargbøl Madsen

## Abstract

The nonselective beta blocker, propranolol, which for decades has been prescribed for treatment of cardiovascular conditions, has recently been used successfully to treat metastatic angiosarcoma. These results have led to an orphan drug designation by the European Medicines Agency for the treatment of soft tissue sarcomas. The anti-tumor effects of propranolol are suggested to involve the reduction of cancer cell proliferation as well as angiogenesis.

Here, we have investigated the anti-angiogenic properties of propranolol in the context of stimulating an anti-tumor immune response. We show that oral administration of propranolol delays tumor progression of MCA205 fibrosarcoma tumors and increases the survival rate of tumor bearing mice. Propranolol works by reducing tumor angiogenesis and facilitating an anti-tumoral microenvironment with increased T cell infiltration and reduced infiltration of myeloid-derived suppressor cells (MDSCs). Using T cell deficient mice, we demonstrate that the full anti-tumor effect of propranolol requires the presence of T cells. Flow cytometry-based analysis and RNA sequencing of FACS-sorted cells show that propranolol-treatment leads to an upregulation of PD-L1 on tumor-associated macrophages (TAMs) and changes in their chemokine expression profile. Lastly, we observe that the co-administration of propranolol significantly enhances the efficacy of anti-CTLA4 therapy.

Our results identify propranolol as an immune modulating agent, which can improve immune checkpoint inhibitor therapies in soft tissue sarcoma patients and potentially in other cancers.

## Introduction

Soft tissue sarcoma (STS) is a heterogeneous group of rare mesenchymal tumors that represent approximately 1% of all adult malignancies. The main treatment option for most patients with localized disease is surgical resection, followed by radiotherapy and chemotherapy in selected subtypes (1). Patients with metastatic diseases have poor clinical outcomes, with an overall survival rate of approximately 12-18 months (2). Over the past decade, the development of immune checkpoint inhibitors (ICIs), which aim to boost anti-tumor T cell activity, has offered new treatment options for patients with melanoma and epithelial cancer, either as a single agent or in combination with conventional therapies (3,4). However, ICIs have achieved suboptimal results on STS patients in two prospective clinical trials (SARC028 and Alliance A091401) of anti-PD1 alone, or in combination with anti-CTLA4 (5,6). Targeting angiogenesis with the tyrosine kinase inhibitor axitinib in combination with anti-PD1 has shown promising preliminary results in selected sarcoma subtypes (7), although the combination has potential overlapping toxicity profiles, which may require temporary treatment cessation of axitinib (8). These studies prompt the combination of ICIs with other less toxic immune modulatory agents to enhance the efficacy.

In recent years, beta adrenergic signaling has been identified as a contributor to tumorigenesis, tumor progression, and metastasis (9–11). The sympathetic nervous system maintains basal concentrations of circulating catecholamines, including epinephrine (Epi) and norepinephrine (NE), which signal through adrenergic receptors (12). The levels of Epi and NE elevate in response to intermittent or chronic stress stimuli, and during tumor progression, neurons from surrounding normal tissues can be recruited to the tumor, further increasing the intratumoral NE level (13,14). Beta adrenergic receptors (ADRBs) are widely expressed in normal tissue. Literature has demonstrated that ADRBs are overexpressed in multiple cancer types such as colon, lung, and breast cancer, compared to normal tissue (15). In the tumor microenvironment (TME), cancer cells as well as immune cells frequently express ADRBs (16). Beta blockers are a class of drugs that competitively blocks the ADRBs and thereby reduces beta adrenergic signaling. Propranolol, a non-selective beta blocker, was first approved in 1967 for cardiovascular indications. However, over the last 50 years, it has been approved for the management of a wide range of other indications, such as essential tremor, prophylaxis of migraines, and benign vascular tumors in infants (17,18). Propranolol treatment has shown to be safe in both adult and pediatric cases (19,20), with few mild side effects, including gastrointestinal disturbances, dizziness, fatigue, and hypotension. The side effects can be effectively managed without the discontinuation of medication (21).

Retrospective studies have shown a correlation between beta blocker usage and reduced overall cancer-associated mortality among cancer patients (22). Combination of propranolol with conventional cancer therapies such as chemotherapy has produced striking responses in hard-to-treat cases such as metastatic angiosarcoma (23–25). The clinical effectiveness of propranolol against angiosarcoma has resulted in an Orphan Drug Designation by the European Medicines Agency (EMA) for the use against STS. Despite promising results in retrospective studies and small-scale clinical studies, the mechanism of action of propranolol in STS is still not completely understood. In angiosarcoma models of immune-deficient mice, propranolol has been shown to reduce cancer cell proliferation and tumor angiogenesis (26). However, solid tumors consist of not only cancer cells, but also immune cells, forming a complex and dynamic TME (27,28). The effect of ADRB stimulation on immune cells in the context of cancer is not well established. While some studies have shown that acute ADRB activation may improve immune function through natural killer (NK) cell mobilization (29), others have observed impaired functions of T cells in the presence of ADRB stimulation (30). In a preclinical study, ADRB activation increased the generation of myeloid derived suppressor cells (MDSCs) *in vitro*, and enhanced their immunosuppressive functions in murine breast cancer models, whereas blockade of ADRB signaling by propranolol reduced the number of MDSCs in both spleen and tumor of tumor bearing mice (31). In clinical settings, propranolol has been shown to reduce surgically induced elevation in peripheral regulatory T cells (Tregs) in breast cancer patients undergoing radical mastectomy (32).

In this study, we investigated how ADRB signaling could alter the composition of the TME in STS using a syngeneic mouse fibrosarcoma model. We demonstrate that propranolol, as a single agent treatment, reduces tumor angiogenesis and improves the anti-tumor response by increasing tumor infiltrating T cells. In addition, propranolol potently reduces intratumoral MDSCs and alters the expression profile of tumor-associated macrophages (TAMs). Finally, we show that propranolol enhances the response to anti-CTLA4 treatment. Our data strongly suggest that ADRB signaling contributes to the formation of an immunosuppressive TME in STS and identifies propranolol as an immune modulating agent that could increase the efficacy of checkpoint inhibitor therapy.

## Methods

### Mouse studies

Animal experiments were conducted at the animal facility of the Department of Oncology, Herlev Hospital, Denmark, under license issued by the Animal Experiments Inspectorate. Daily maintenance of C57BL/6 and NMRI-nu/nu mouse stocks were performed by the animal caretakers at the animal facility. Experimental mice were females between 8 and 16 weeks of age. The right flank of mice was subcutaneously (s.c.) injected with 1 × 10^6^ MCA205 cells. Tumor dimensions (length and width) were measured three times a week with a digital caliper. Tumor volumes were calculated by the formula: 0.5 x length x width^2^. Mice were weighed weekly and inspected visually three times a week to ensure wellbeing. The experimental endpoint was defined as tumor volume exceeding 1000 mm^3^. The humane endpoint was defined as more than 20% weight loss, ulcerating tumors, or wounds on tumors of more than 5 mm in length. Mice were sacrificed by cervical dislocation and the tumors were collected for further analysis unless otherwise noted.

### Cell lines

The MCA205 cell line was purchased from Merck (SCC173) and cultured in complete growth medium, consisting of RPMI-1640 GlutaMAX™ Supplement HEPES, 20% FBS, 1% sodium pyruvate, 1% nonessential amino acids, and 1% P/S (All from Gibco). The complete growth medium was supplemented with β-mercaptoethanol (Gibco) at a concentration of 55µM before use. The SVR cell line was purchased from ATCC (CRL-2280) and cultured in complete growth medium, consisting of DMEM, 10% FBS, and 1% P/S (All from Gibco). The MC38 cell line was retrieved from cell line biobank in the group and cultured in complete growth medium, consisting of DMEM, 10% FBS, and 1% P/S (All from Gibco).

### Therapies

(±) Propranolol hydrochloride (propranolol) (Sigma) was dissolved in room temperature drinking water at 0.5 g/L and offered to mice ad libitum, starting on the day of cancer cell inoculation. Water bottles for all mice were changed every 3-4 days. In checkpoint inhibitor treated groups, mice were treated intraperitoneally (i.p.) three times a week, starting one day after cancer cell inoculation, with anti-PDL1 mAb (200 µg/mouse; clone 10F.9G2, BioXcell), anti-CTLA4 mAb (200 µg/mouse; clone 9D9, BioXcell), anti-PD1 mAb (200 µg/mouse; clone RMP1-14, BioXcell), alone or in combinations until endpoint or one week after tumor clearance. Control mice were treated with PBS following the same treatment scheme.

### Bone marrow derived macrophages (BMDMs)

BMDMs were generated from the marrow of femurs and tibias from female C57BL/6 mice. Bones were rinsed in 70% ethanol and in PBS, and the bone marrow cells were flushed with PBS using a 25G needle. After red blood cell lysis with RBC lysis buffer (Qiagen), cells were cultured in BMDM growth media, consisting of Iscove’s modified Dulbecco medium (IMDM, Sigma Aldrich), supplemented with 20% FBS, 1% P/S, and 20 ng/mL recombinant human M-CSF (R and D system), at a concentration of 1 × 10^6^ cells/mL. The growth medium was refreshed on day 3. Differentiated macrophages were collected on day 7, and the purity determined to be more than 98% by multicolor flow cytometry, using cell surface markers CD11b and F4/80. The macrophages were cultured under different conditions for 24 hours for specific macrophage polarization. For M1 polarization of macrophages, 100 ng/mL LPS (Invitrogen) and 40 ng/mL IFNγ (PeproTech) were used. For M2 polarizations, 40 ng/mL IL4 (PeproTech) was used. For adrenergic stimulation and blockade, (±) isoprenaline hydrochloride (isoprenaline) (Sigma Aldrich) or propranolol was dissolved in BMDM growth media and added to the cells.

### Flow cytometry analyses

Excised tumors were minced with scissors and enzymatically digested in RPMI medium containing 2.1 mg/ml collagenase type 1 (Worthington), 75 µg/ml DNase I (Worthington), 5 mM CaCl_2_, and 1% P/S. Cells were then filtered through a 70 µm cell strainer. After red blood cell lysis, cells were incubated with FcR block (Miltenyi) for 10 minutes at 4°C. Dead cells were excluded using Zombie Aqua viability dye (BioLegend). For membrane staining, cells were incubated with antibody mix at 4°C in the dark for 20 minutes.

For intracellular IFNγ and TNFα staining, isolated splenocytes from tumor free mice were first stimulated with in vitro cultured MCA205 cancer cells in a 24-well plate for 24 hours in MCA205 complete growth media. A ratio of 2 × 10^6^ splenocytes to 200,000 cancer cells were used. GolgiPlug (1/1000, BD Biosciences) was added to the mixed culture and incubated for an additional 4 hours. Non-adherent cells were collected and stained for surface markers before permeabilization using a FoxP3/transcription factor staining buffer set (eBioscience), according to manufacturer’s instruction. The cells were stained with anti-IFNγ and anti-TNFα (Biolegend) at 4°C dark for 35 minutes. All antibodies used are listed in the supplementary table 1. Data were acquired on a FACSCanto II (BD) or Quanteon (NovoCyte) flow cytometer after appropriate compensation using single-stained cells or compensation beads (BD Biosciences). Data were analyzed with Flow Jo software (Tree Star).

### Gene expression analyses

Excised MCA205 tumors were stored in RNAlater (Invitrogen) at -80°C until RNA extraction. Tumors were placed in buffer RLT (Qiagen), and mechanically homogenized with Tissue Lyser (Qiagen). Total RNA extraction was performed with the RNeasy Mini kit (Qiagen), following the manufacturer’s instruction. A maximum of 1 µg of RNA, measured by NanoDrop Spectrophotometer (Thermo Scientific), was reverse transcribed into cDNA using iScript cDNA synthesis kit (Biorad), according to manufacturer’s protocol. qRT-PCR was performed using the Brilliant III Ultra-Fast SYBR® dye system (Agilent) with ROX as a reference dye. The loaded plates were run on an AriaMX Real-Time PCR System with the thermal profile: 1 cycle at 95°C for 3 min, followed by 40 cycles of 95°C for 5 sec, 60°C for 20 sec. This was followed by a melting curve analysis of 65–95 °C with 0.5°C increment, 5 s per step. Quantitative qRT-PCR data were normalized to the expression level of the housekeeping gene *Actb*. Data were analyzed using the 2^−ΔΔCt^ method, and fold change of treated group compared to control group was calculated. Primer sequences are listed in supplementary table 2.

### Cell sorting and RNA sequencing

For TAM isolation, single cell suspensions of MCA205 tumors from control and propranolol treated mice were labeled with CD45 microbeads (Miltenyi) and the CD45+ cells were purified using magnetic separation. The CD45+ cell fractions were stained with Zombie Aqua viability dye, anti-CD11b, and anti-F4/80 antibodies. CD11b+ F4/80+ live cells were sorted by FACS on the ARIA III (BD Biosciences). Purity of samples was tested to be at least 96%. RNA from sorted TAMs were extracted immediately after sorting, using RNeasy Mini Kit (Qiagen) according to the manufacturer’s instructions. Quality of RNA samples was determined by Bioanalyzer (Agilent). Only samples with RNA integrity number (RIN) of more than 7.5 were used for RNA sequencing.

### RNA sequencing

In total 400 ng RNA was prepared for sequencing using polydT enrichment according to the manufacturer’s instructions (Illumina). Library preparation was performed using the NEBNext RNA library prep kit for Illumina. The library quality was assessed using a Fragment Analyzer followed by library quantification using the Illumina library quantification kit. Libraries were sequenced on a NovaSeq 6000 platform (Illumina) to a minimum depth of 30 million reads per sample. Sequenced reads were aligned to the reference mm10 genome for mouse using STAR, version 2.7.1 (33). For quality control FastQC version 0.11.8 (34), AdapterRemoval version 2.3.0 (35), FastQ Screen version 0.13.0 (36), SAMtools version 1.9 (37,38), Picard Toolkit (39), Qualimap version 2.2.2 (40), and RNA-SeQC version 2.3.4 (41) were used. MultiQC version 1.9 (42) was used to assess the quality metrics. High-quality reads were filtered by MAPQ scores (MAPQ > 30) using SAMtools. Mates unmapped and secondary alignments were removed with samtools view -F 780. The gene expression count matrix was generated using featureCounts version 1.6.4 (43) and GENCODE gene annotation with the parameters “-p -t exon”.

### Analysis of RNA sequencing

For all analysis of the RNA seq data R version 4.0.3 was used. The DESeq2 package, version 1.30 (12), was acquired for the analysis of differentially expressed genes (Cut off p-adjusted value <0.05 and log2 fold change >1.5). To evaluate the expression all differentially expressed genes and selected genes of interest, heatmaps were made using the package Pheatmap version 1.0.12 (44) with parameters clustering_method = “ward.D” and scale = “row”, i.e. z-score–DESeq normalized counts. For visualization of specific genes of interest, the ggplot2 package version 3.3.3 was used. Reads were normalized to transcript per kilobase million (TPM). For the GoSeq analysis, compareCluster from the clusterProfiler package version 3.18.0 was used. Redundant GOterms were excluded using the Simplify command. The Benjamini-Hochberg method (45) and p-adjusted-value cut-off <0.05 were set as criterion for the GOSeq analysis. The ggplot2 package version 3.3.3, was used to visualize the top ten GOterms.

### Immunohistochemistry

MCA205 tumors were fixed in 4% formaldehyde using Sarstedt formalin system overnight at 4 °C. Samples were transferred to 70% ethanol and stored at 4 °C until paraffin embedding. Tissues were embedded in paraffin and cut into 4 µm tissue sections. Hematoxylin and eosin staining was performed according to standard protocols. For CD34 immunostaining, 10 mM citrate buffer, Ph antigen retrieval, and rat anti-mouse CD34 (1:400, MEC 14.7, Novus Biologicals) were used. For CD8 immunostaining, proteinase K buffer antigen retrieval, rat anti-mouse CD8a (1:50, 4SM16, Invitrogen) were used. For Ki67 immunostaining, Target Retrieval Solution, Citrate pH (DAKO), rabbit anti-mouse Ki67 (1:100, SP6, Abcam) were used. The sections were then stained with polyclonal rabbit anti-rat IgG (1:200, Dako), or EnVision rabbit (Dako) with NOVAred substrate (Vector). The tissue sections were counterstained with hematoxylin. The percentage of CD8+ or Ki67+ cells were quantified with Qupath software (46), using positive cell detection with optimized parameters. The percentage of CD34+ areas were quantified with Qupath software (ver. 0.2.3), using trained positive pixel classifier.

### Statistical analyses

Data analyses and graph generations were performed with Prism 6 (GraphPad) unless otherwise stated. Statistical analyses were performed using multiple t test with Bonferroni-Dunn correction for multiple comparisons. Tumor growth related statistics were analyzed with the open-access web tool TumGrowth (kroemerlab.shinyapps.io/TumGrowth/) (47). Default settings were used.

## Results

### Blockade of ADRB signaling delays sarcoma growth

To investigate the importance of ADRB signaling for sarcoma growth, we evaluated the effect of the non-selective ADRB blocker propranolol on a murine model of fibrosarcoma. C57BL/6 mice were inoculated s.c. with MCA205 fibrosarcoma cells and propranolol was administered orally via the drinking water. Propranolol treatment resulted in delayed tumor growth and increased median survival rate of the mice from 18 days to 21 days (fig 1A and B). To confirm the anti-tumor effect of propranolol, all tumors were excised and weighed on day 16 in a separate experiment. The weight of tumors from propranolol treated mice were significantly lower than control mice (fig 1C). Since propranolol has been reported to have anti-proliferative effects on a variety of epithelium-derived cancer cells (48), we investigated if the delay in tumor growth could be due to a direct inhibition of MCA205 proliferation. We first confirmed the gene expression of *Adrb1* and *Adrb2* in *in vitro* cultured MCA205 cells by qRT-PCR (fig 1D). By subjecting cultured MCA205 cancer cells to propranolol in concentrations ranging from 0.4 µM to 200 µM, we observed a dose dependent inhibition of MCA205 proliferation and reduction of viability *in vitro* (fig 1E and F). A similar effect was observed on the murine angiosarcoma cell line SVR and murine colon adenocarcinoma cell line MC38 (supp. fig. S1). To examine if this anti-proliferative effect could be observed *in vivo*, we stained tumor sections from control and propranolol groups for the proliferation marker Ki67 by immunohistochemistry. Surprisingly, the expression levels of Ki67 in tumors from both groups were similar, both in the tumor core and at the invasive front (fig 1G and H). This indicated that the effect of propranolol on tumor growth was unlikely due to direct inhibition of cancer cell proliferation.

**Figure 1.**
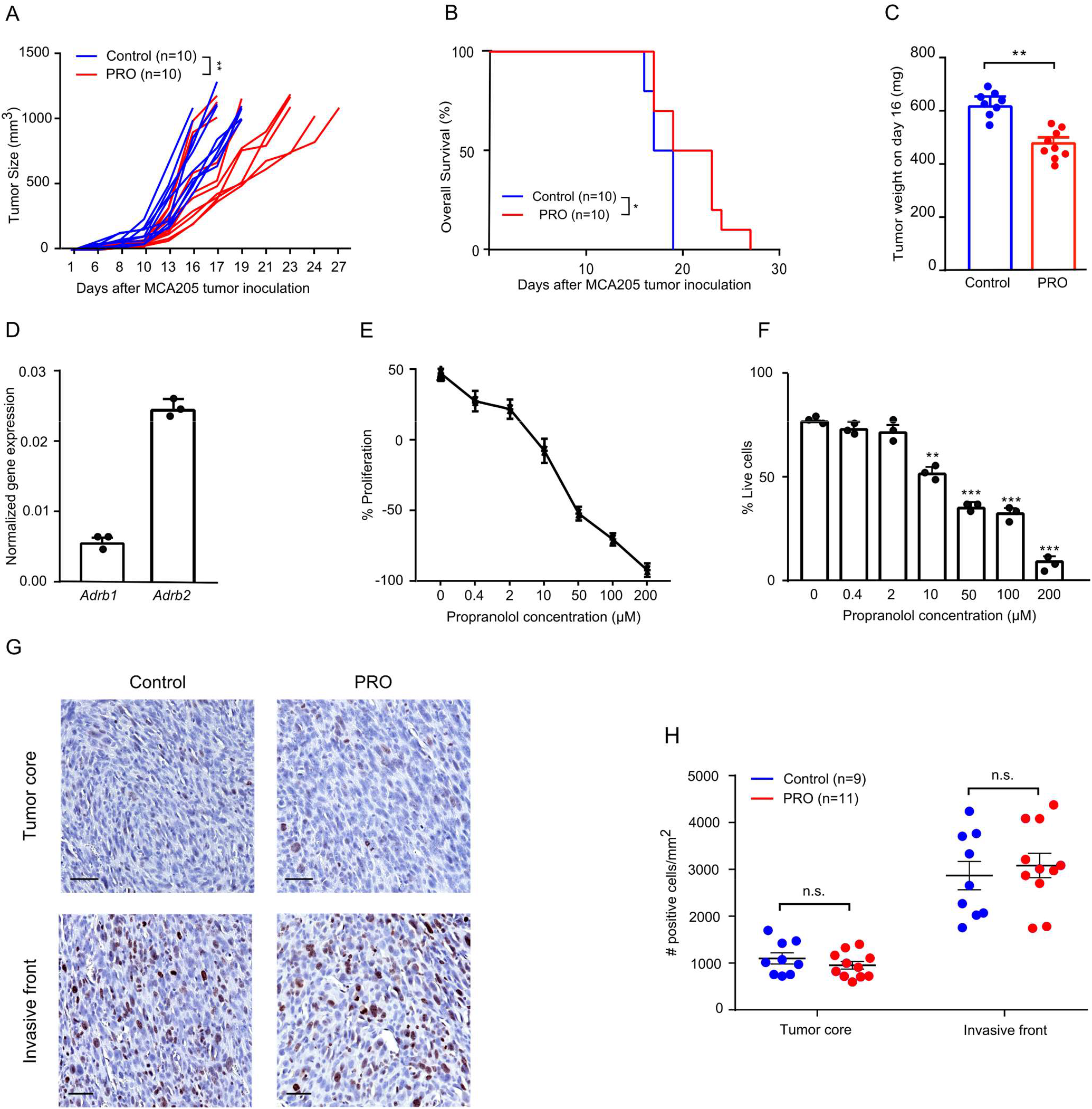
Pharmacological ADRB blockade by propranolol delays tumor growth of MCA205 fibrosarcoma. **A and B**, Tumor growth curves (A) and survival rates (B) of control, and propranolol (PRO) treated mice (n=10). **C**, Tumor weight on day 16 (n=8-9). **D**, Quantification of gene expression of *Adrb1, Adrb2* in *in vitro* cultured MCA205 cancer cells by qRT-PCR (normalized to *Actb*) (n=3, with 3 technical replicates each). **E and F**, Proliferation and viability of MCA205 cells upon propranolol treatment were quantified by cell counting with hemocytometer. Dead cells were identified using trypan blue. Proliferation (**E**) and viability (**F**) of MCA205 cells after 24 hours of propranolol treatment at various concentrations, compared to pre-treatment (n=3). **G**, Representative Ki67 (brown) IHC staining on MCA205 tumor tissues from control and propranolol treated mice; hematoxylin counterstain; scale bar 50µm. **H**, Dot plot showing quantification of Ki67 staining in 9 control mice and 11 propranolol treated mice. **p<0.01, ***p<0.005, n.s. not significant, according to multiple t test with Bonferroni correction. Tumor growth and survival data were analyzed using TumGrowth software. Mean ± SEM are depicted for C&H; mean ± SD are depicted for D-F.

Propranolol has also been demonstrated to be a potent anti-angiogenic agent in preclinical studies by reducing the expression of vascular endothelial growth factor A (VEGFA) (18). Therefore, we measured *Vegfa* gene expression in whole tumor RNA from control and propranolol treated mice using qRT-PCR. Propranolol treatment resulted in a 50% reduction in intratumoral *Vegfa* gene expression (fig 2A), while having no effect on expression of the *Kdr* gene encoding the primary receptor of VEGFA, vascular endothelial growth factor receptor 2 (VEGFR2) (Fig 2B). To confirm the anti-angiogenic effect of propranolol, we immuno-stained paraffin-embedded tissue sections for the endothelial marker CD34 and quantified the CD34-positive area. Tumor sections from the propranolol treated group had a significantly lower CD34-positive area compared to the control group, confirming the anti-angiogenic effect of propranolol (fig 2C and D).

**Figure 2.**
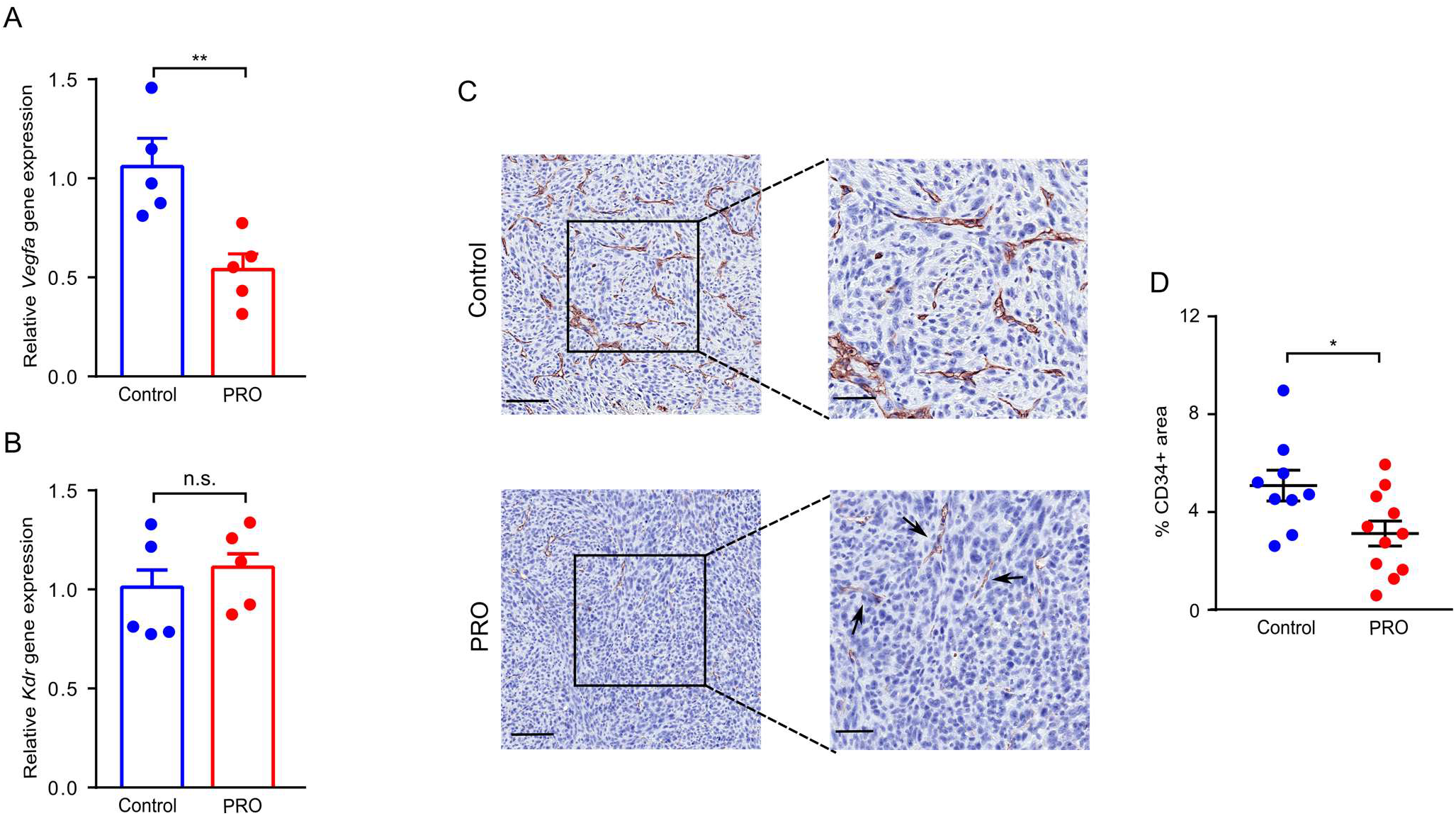
Propranolol treatment reduces MCA205 tumor angiogenesis. **A and B**, Quantification of angiogenic marker Vegfa (A) and Kdr (B) gene expression in whole tumor RNA from control mice and propranolol (PRO) treated mice by qRT-PCR (n=5). **C**, Representative CD34 (brown) IHC staining of MCA205 tumor tissue from control mice or propranolol treated mice; hematoxylin counterstain; scale bar 100µm (left panels), and 50µm (right panels). **D**, Dot plot showing quantification of CD34 staining by percentages of DAB+ area in 9 control mice and 11 propranolol treated mice. *p<0.05, **p<0.01 according to multiple t test with Bonferroni correction. Mean ± SEM are depicted.

### The anti-tumor effect of propranolol involves T cell activity

To investigate if the mechanisms of action of propranolol extended beyond anti-angiogenic effects, we analyzed if propranolol affected the infiltration of T cells. The cellular composition of the lymphoid compartment of the MCA205 tumors was analyzed by multicolor flow cytometry. Propranolol led to an increased infiltration of CD4+ T cells into the tumors, without altering the ratio of Tregs to total CD4+ T cells (Fig 3A-C). There was also a trend towards an increase in intratumoral CD8+ T cells in the propranolol treated mice (Fig 3D). The expression levels of the T cell activation marker CD137 and exhaustion marker PD1 on CD4+ and CD8+ T cells were similar between the control and propranolol group (supp. fig. S2A). We did not observe changes in the abundance of NK cells in the tumors (supp. fig. S2B).

**Figure 3.**
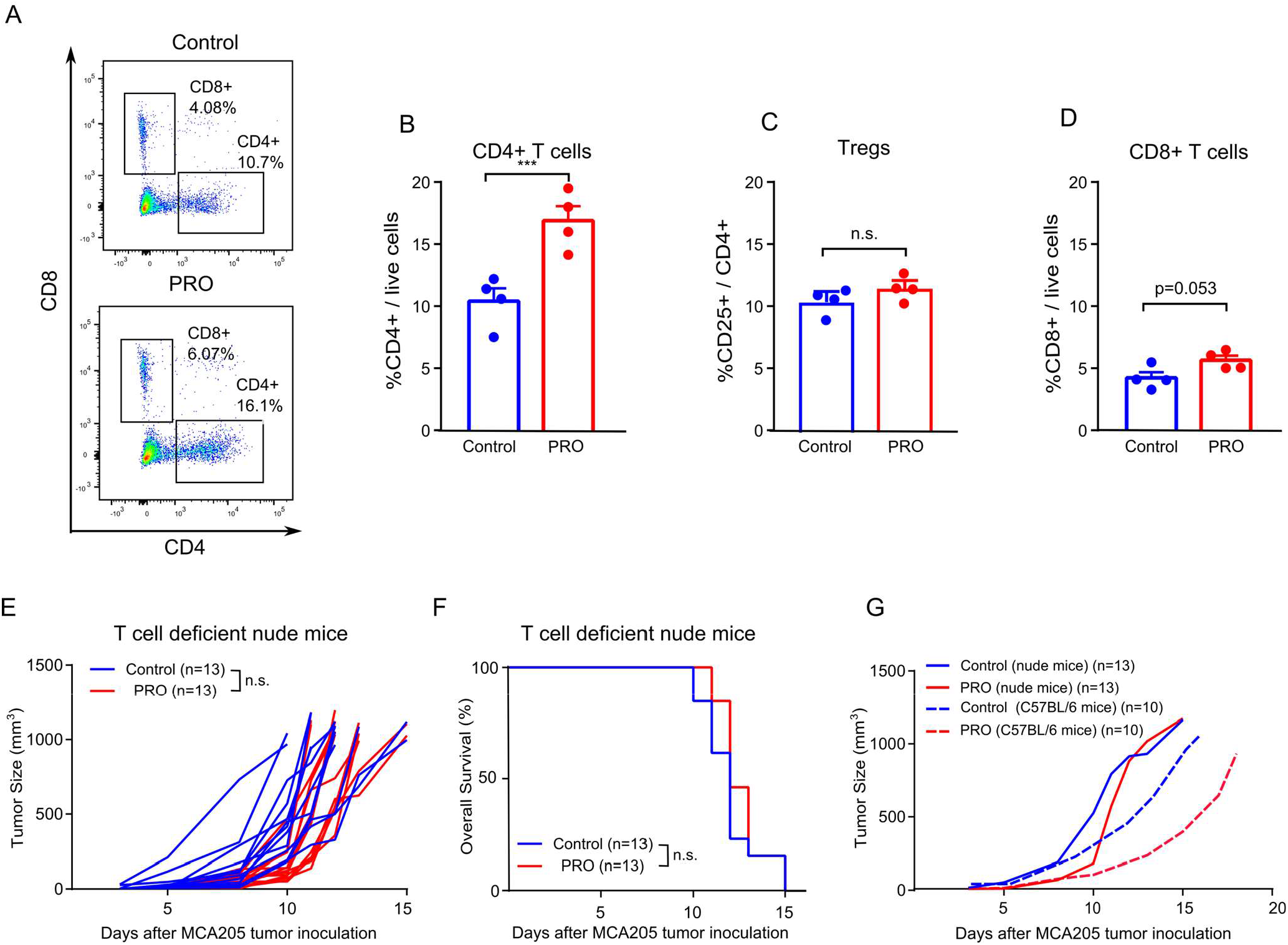
Propranolol treatment_increases CD4+ T cell infiltration in MCA205 tumors. **A-D**, Single cell suspensions were made from excised MCA205 tumors at the experimental endpoint and analyzed by flow cytometry. **A**, Representative flow cytometry dot plot of CD4+ and CD8+ T cells in the TME. Quantification of CD4+ T cells **(B)**, regulatory T cells (**C**), and CD8+ T cells (**D**) (n=3-5). **E-F**, Tumor growth kinetics **(E)** and survival rates **(F)** of control, and propranolol-treated (PRO) nude mice (n=13). **G**, Comparison of average tumor growth between nude mice and immune competent mice treated with propranolol. For tumor growth and Kaplan–Meier curves, statistical analyses were performed using TumGrowth software. For other comparisons, multiple t tests with Bonferroni correction for multiple comparison were used. *p<0.05, **p<0.01, ***p<0.005, n.s.: not significant. Mean ± SEM are depicted.

To directly test if the therapeutic efficacy of propranolol is dependent on the presence of T cells, we inoculated T cell deficient nude mice with MCA205 cells and treated them with propranolol. A similar experiment with immune-competent mice was conducted concurrently. In nude mice, the growth rate of MCA205 tumors and survival rate of mice were comparable between control and propranolol-treated mice (Fig 3E and F). In immune-competent mice, the difference between the tumor growth rate of propranolol treated and control mice was apparent (Fig 3G). These results suggested that the anti-tumor effect of propranolol depends on the presence of T cells.

### Blockade of ADRB signaling affects the myeloid compartment of the TME

Since immunosuppressive cells of the myeloid lineage can contribute to tumor progression in a variety of cancer types (49), we investigated if propranolol affected the cellular composition of the myeloid compartment in the TME of MCA205 tumors. Flow cytometry-based analysis of the TME showed that propranolol treatment led to a striking reduction in the abundance of intratumoral myeloid-derived suppressor cells (MDSCs) (Fig 4A and B). No difference was observed in the numbers of tumor-associated macrophages (TAMs) (Fig 4C). TAMs are heterogeneous cells, which depending on the context, can acquire an immunosuppressive phenotype (50). We measured the surface expression of MHC II, PDL1, and CD206 on TAMs and found an increase in PDL1 expression, and a trend towards increased MHC II expression on TAMs from propranolol treated mice (fig 4F). There was no difference in the expression level of the M2 marker CD206 (Fig 4G). In nude mice, we also observed an increase in PDL1 expression on TAMs in MCA205 tumors from propranolol treated mice (fig 4E), although less pronounced than in the presence of T cells (compare Fig. 4D and Fig. 4E). This suggests that macrophages can be affected directly by the blockade of ADRB.

**Figure 4.**
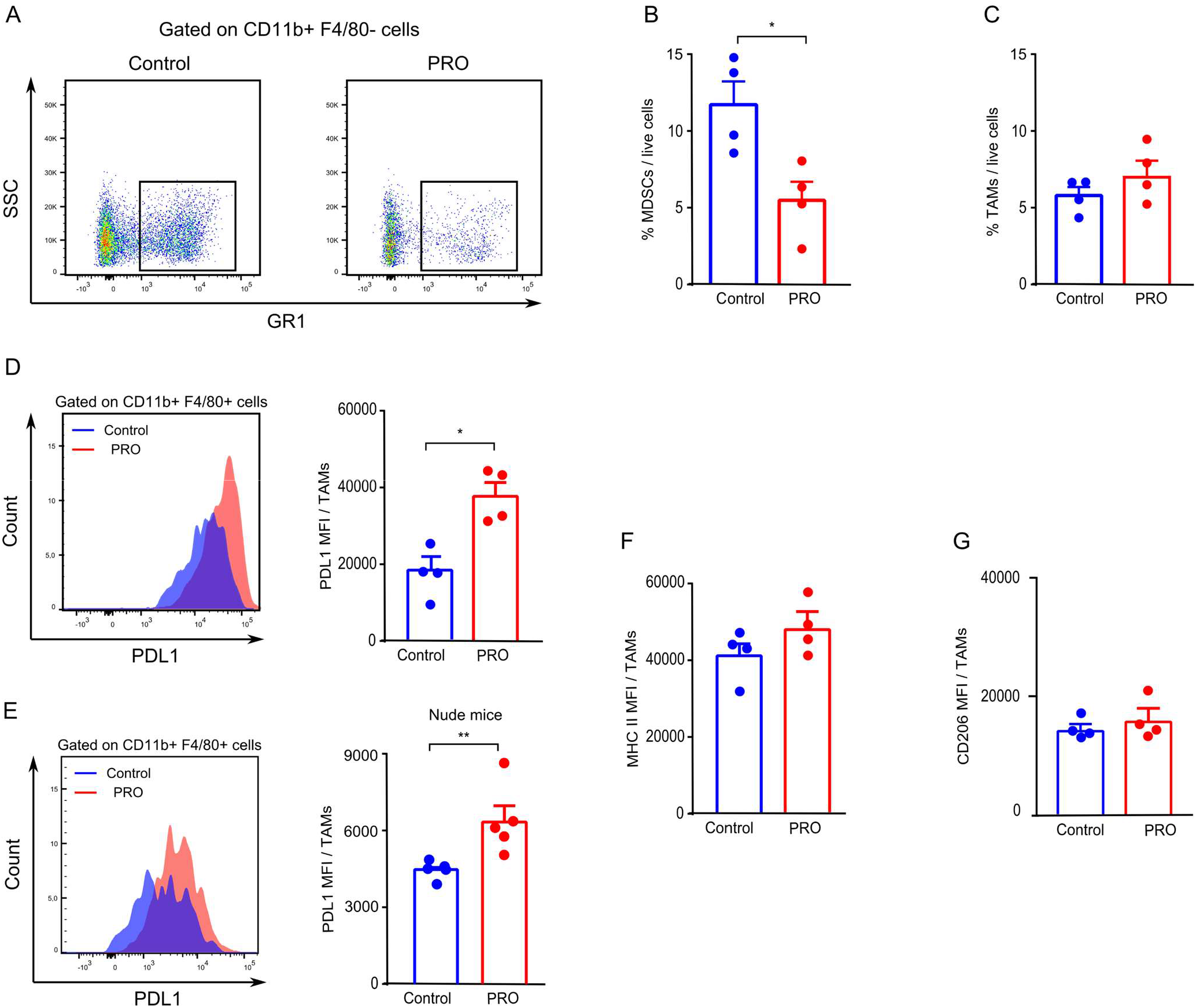
ADRB blockade by propranolol reduces intratumoral MDSCs and upregulates PDL1 expression on tumor associated macrophages (TAMs). Single cell suspensions were made from excised MCA205 tumors in immune competent mice and T cell deficient nude mice at the experimental endpoint and analyzed by flow cytometry (n=3-5). **A and B**, Representative flow cytometry dot plot **(A)** and quantification of MDSCs (CD11b+ F4/80-GR1+) **(B)** in the TME of tumors from control and propranolol (PRO) groups in immune competent mice. **C**, Quantification of TAMs (CD11b+ F4/80+) in samples from control and propranolol-treated immune competent mice. **D and E**, Representative histograms and quantifications of mean fluorescence intensity (MFI) of PDL1 on TAMs from immune competent mice (**D**), and T cell deficient nude mice (**E**). MFI quantification of MHC II (**F**) and CD206 (**G**) expression on TAMs in immune competent mice. Multiple t tests with Bonferroni correction for multiple comparison were used for statistical testing. *p<0.05, **p<0.01. Mean ± SEM are depicted.

### ADRB signaling affects the phenotype of macrophages *in vitro*

To test how macrophages respond to ADRB stimulation, we generated BMDMs. The differentiated macrophages were treated with the pan-ADRB agonist isoprenaline or with propranolol. The expression of PDL1 was increased by propranolol treatment while isoprenaline had no effect (fig 5A). We then evaluated the expression of the M1 markers CD86 and MHC II, and the M2 marker CD206 upon ADRB stimulation or subsequent blockade under M1 or M2 polarizing conditions. Isoprenaline dampened BMDMs’ response to IFNγ and LPS stimulation, demonstrated by a reduction in MHC II and CD86 expression. Subsequent blocking of ADRB with propranolol rescued the expression of these two markers (fig 5B). In the presence of the M2 polarizing cytokine IL4, isoprenaline enhanced the expression of CD206, skewing BMDMs further towards M2-like polarization. Blockade of ADRB by propranolol neutralized the effect (fig 5C). Interestingly, propranolol alone dampened the M2 polarizing effect of IL4; the expression levels of MHC II, CD86, PDL1, and CD206 on IL4 and propranolol treated BMDMs were similar to those of untreated control cells (fig 5C, and supp. fig. S3A-C).

**Figure 5.**
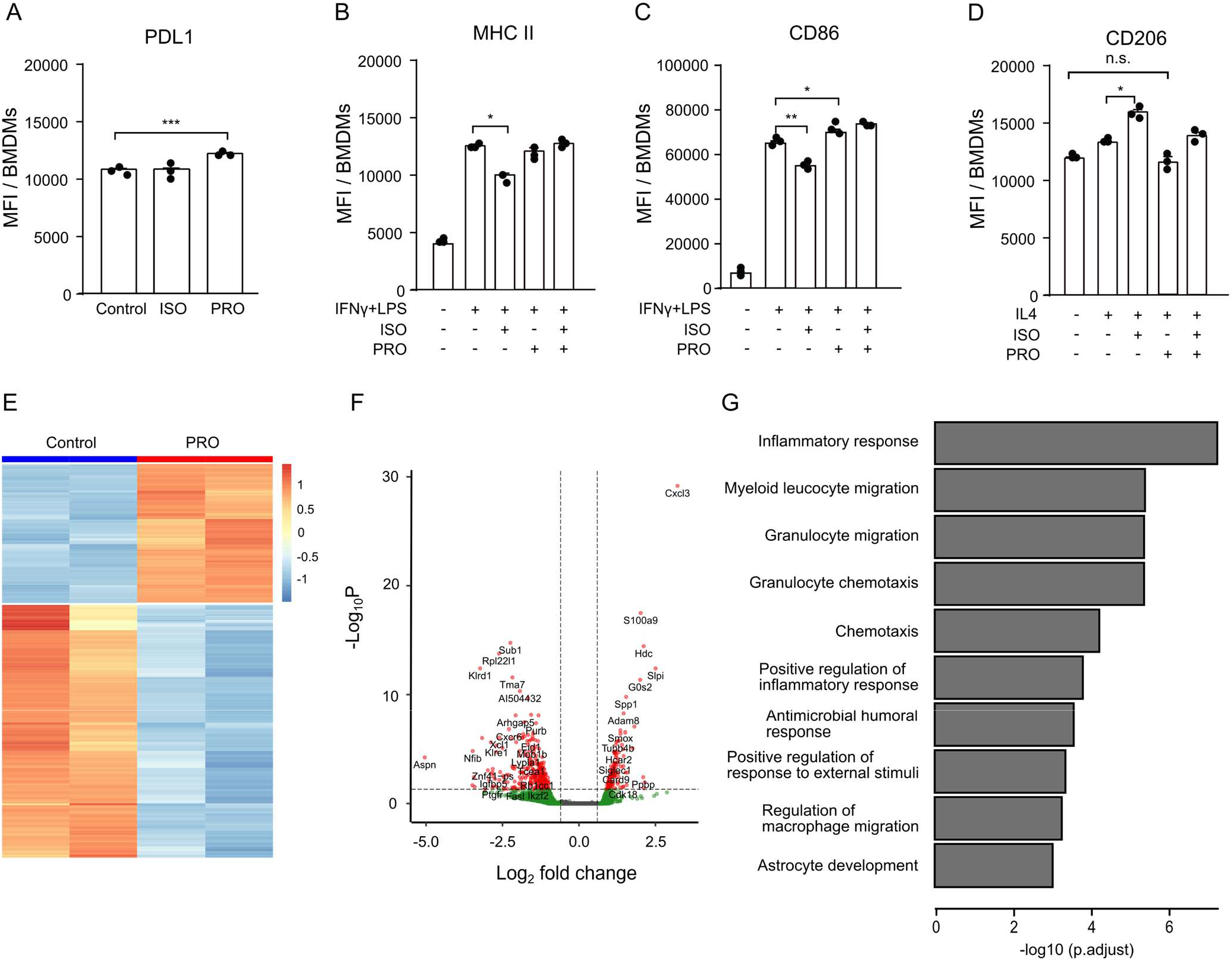
Macrophages are sensitive to ADRB stimulation and propranolol treatment leads to TAMs with distinct gene expression profiles. **A**, Quantification of PDL1 MFI on non-polarized macrophages upon isoprenaline (ISO) or propranolol (PRO) treatment. isoprenaline and/or propranolol’s effect on the surface expression of MHC II (**B**), CD86 (**C**), or CD206 (**D**) under M1 polarizing condition with IFNγ and LPS or under M2 polarizing condition with IL4. **E and F**, RNA-seq analysis showing heatmap (**E**) and volcano plot (**F**) of genes differentially regulated in TAMs from control or propranolol treated mice. Data show two biological replicates. **G** Gene ontology analysis showing the biological processes most significantly enriched within genes that are differentially expressed between TAMs isolated from control mice or propranolol treated mice. *p<0.05, **p<0.01, ***p<0.005, according to multiple t test with Bonferroni correction for multiple comparison. Mean ± SD are depicted.

### TAMs from MCA205 tumors of propranolol treated mice have a distinct gene expression profile

*Ex vivo* generated BMDMs are responsive to ADRB signaling and blockade. To examine how propranolol affects the phenotype of TAMs *in vivo*, we FACS-sorted TAMs from MCA205 tumors of propranolol treated or control mice and extracted RNA for RNA sequencing (RNAseq). TAMs from propranolol treated mice had a distinct gene expression profile compared to TAMs from control mice (fig 5E). We identified 390 genes, which were regulated by more than 1.5 fold (p<0.05) in response to propranolol treatment (fig 5F). Top ten most up-regulated and down-regulated genes are listed in table 1 and the complete list of differentially regulated genes can be found in supplementary table 3. Gene ontology enrichment analysis revealed that the differentially regulated genes were involved in biological processes including inflammatory response, myeloid leukocyte migration, and granulocyte migration (fig 5G).

**Table 1.**
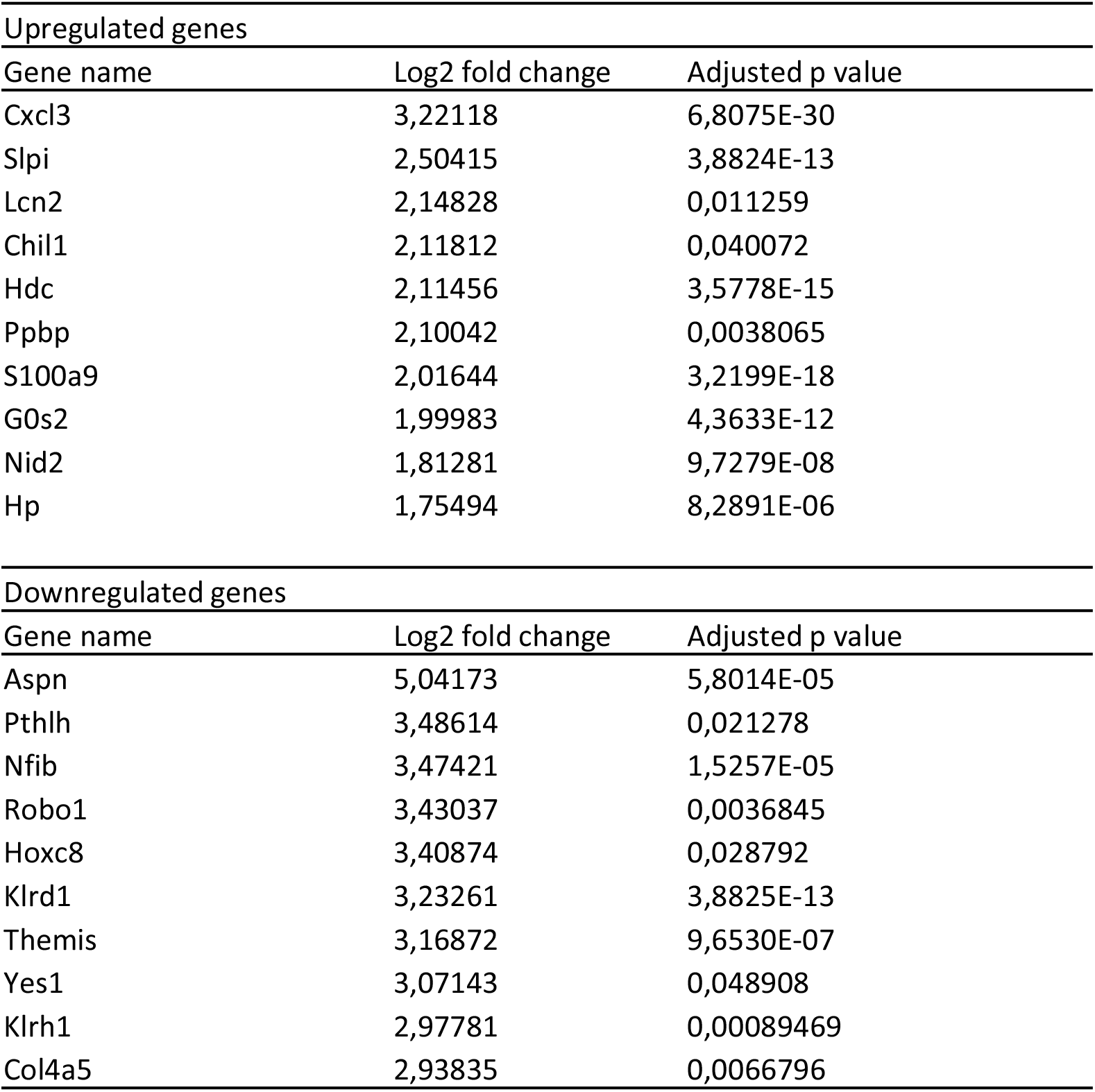
Top 10 most upregulated and down regulated genes

### Propranolol enhances the effect of anti-CTLA4 treatment

A high abundance of tumor-infiltrating T cells is a predictive marker of ICI efficacy (51), and a low number of MDSCs has been shown to correlate with an increased chance of responding to anti-CTLA4 therapy (52). Propranolol treatment led to increased T cell infiltration and reduced intratumoral MDSCs in MCA205 tumors and we therefore decided to combine propranolol with ICI therapy (fig 6A). First, propranolol was combined with anti-PD1 treatment. MCA205 tumors were minimally responsive to anti-PD1 treatment, and the tumor growth and survival rate of mice were comparable between the two ICI treated groups with or without propranolol (fig 6B-D). Similar results were obtained when propranolol was combined with anti-PDL1 treatment (supp. fig. S4A), suggesting that ADRB blockade did not affect therapy targeting PD1/L1 axis in this tumor model. Next, we tested the effect of anti-CTLA4 in combination with propranolol. Anti-CTLA4 alone led to temporary stabilization of MCA205 tumors, before the tumors became resistant to therapy. Interestingly, the addition of propranolol further delayed tumor growth and improved the survival rate compared to anti-CTLA4 alone (fig 6E-G). The treatment combination also improved the response rate to treatment, from 8/15 (53%) in anti-CTLA4 group to 12/15 (80%) in anti-CTLA4+PRO group. One mouse in the anti-CTLA4+PRO group showed complete tumor clearance, while none in the anti-CTLA4 single treated group achieved tumor free status. To gain insight into the mechanism of action of the observed synergistic effect, we performed immunohistochemical staining of CD8 and CD34 on paraffin embedded tissue sections from mice treated with anti-CTLA4 with or without propranolol. Anti-CTLA4 treated mice had a marked increase in tumor infiltrating CD8+ T cells compared to control mice, and the combination of propranolol and anti-CTLA4 led to a further increase in the number of tumor-infiltrating CD8+ T cells (fig 7A and B). The anti-angiogenic effect of propranolol as a single agent treatment was also observed when used in combination with anti-CTLA4 (fig 7C and D).

**Figure 6.**
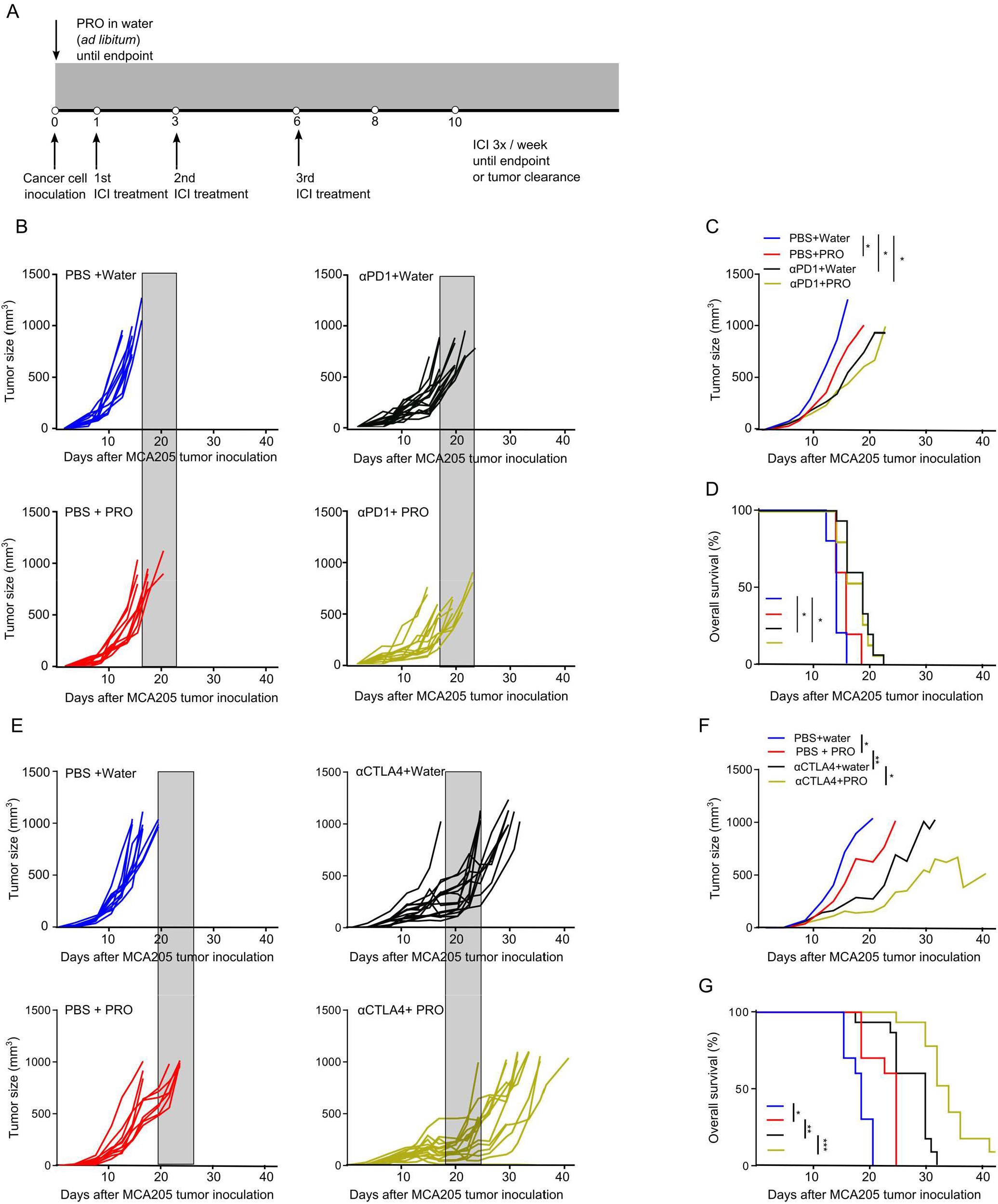
ADRB blockade by propranolol improves the efficacy of anti-CTLA4, but not anti-PD1 treatment in the MCA205 tumor model. **A**, Treatment regimen and experimental setup of immune checkpoint inhibitor (ICI) tumor studies. **B-D**, Tumor growth kinetics (**B**), average tumor sizes per group (**C**), and Kaplan-Meier survival curves (**D**) of C57BL/6 mice inoculated with MCA205 cancer cells, treated with anti-PD1 and propranolol (PRO). **E-G**, Tumor growth kinetics (**E**), average tumor sizes per group (**F**), and Kaplan-Meier survival curves (**G**) of C57BL/6 mice inoculated with MCA205 cancer cells, treated with anti-CTLA4 and propranolol. Each line represents one animal. n=10-15 per group. Shaded areas, included for easy comparison of the different treatment groups, represent the terminal tumor growth time frame of the PBS+water control group. Statistical analyses were performed using TumGrowth software. *p<0.05, **p<0.01, ***p<0.005.

**Figure 7.**
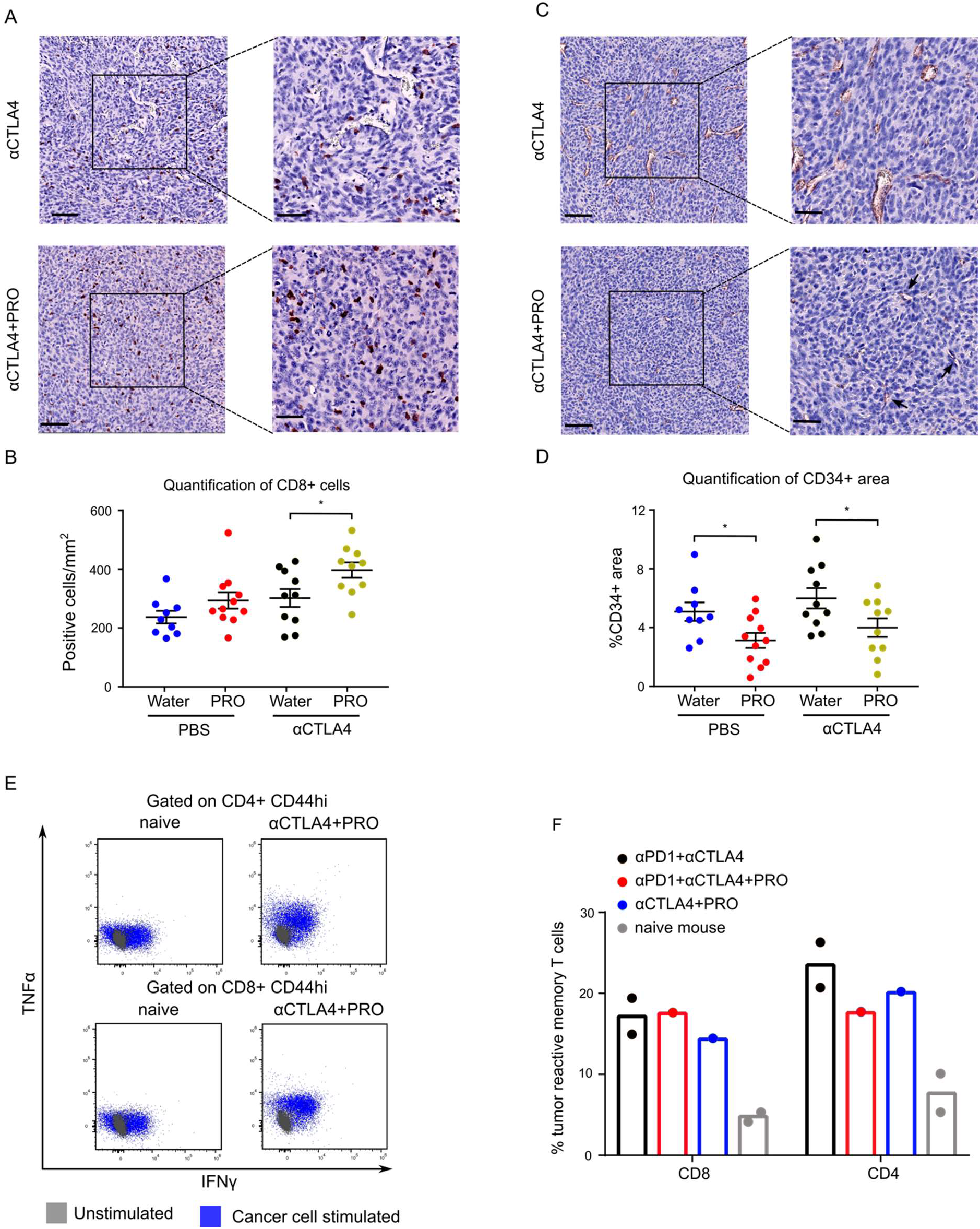
Propranolol combined with anti CTLA4 increases the number of intratumoral CD8+ T cells, reduces tumor angiogenesis, and provides long lasting immune memory against MCA205 cancer cells. **A and C**, Representative CD8 (**A**) and CD34 (**C**) IHC staining of MCA205 tumor tissues; hematoxylin counterstain; scale bar 100µm (left panels), and 50µm (right panels). **B and D**, Dot plot showing quantification of CD8 (**B**), and CD34 staining (**D**). Splenocytes from cured mice were re-stimulated *ex vivo* with MCA205 cancer cells for 24 hours, and intracellular cytokine expressions were quantified by flow cytometry. **E**, Representative flow cytometry dot plots showing the expression of TNFα and IFNγ on CD4+ T cells (upper panels) or CD8+ T cells (lower panels) with memory phenotype from naïve mice (left panels) or anti-CTLA4+PRO treated mice that had shown complete tumor regression(right panels). **F**, Percentages of tumor reactive T cells (IFNy+ or TNFa+) with memory phenotype (CD44hi) among CD8+ or CD4+ T cells after 24-hour ex vivo re-stimulation. *p<0.05 according to multiple t test with Bonferroni correction. Mean ± SEM are depicted.

We then tested if propranolol could increase the response rate of an already very effective treatment regimen of combined anti-PD1 and anti-CTLA4 antibodies. Propranolol did not alter the effectiveness of the maximum dose of ICI treatment (supp. fig S4). In the ICI treated groups with or without propranolol, 13/15 mice responded to treatment, and 3 mice experienced complete tumor regression. Out of the 6 tumor-free mice from both ICI treated groups, 3 mice relapsed after treatment discontinuation.

### Anti-CTLA4 and propranolol treatment can lead to the development of immune memory against MCA205 cancer cells

To test if mice that showed complete tumor regression developed immune memory against the cancer cells, we re-inoculated the mice with MCA205 cancer cells three months after tumor clearance. All mice remained tumor-free during the one-month observation period, while 5 out of 5 tumor naïve mice developed tumors within 6 days of inoculation. We then sought to evaluate the amount of tumor reactive T cells in cured mice, compared to tumor naïve mice. Isolated splenocytes from tumor free mice were stimulated with MCA205 cells and the expression levels of IFNγ and TNFα in T cells were examined by flow cytometry. T cells positive for either IFNγ or TNFα were defined as tumor reactive. Splenocytes from tumor free mice treated with anti-PD1+anti-CTLA4, anti-PD1+anti-CTLA4+PRO, and anti-CTLA4+PRO harbored higher number of tumor reactive T cells with a memory phenotype against MCA205 cancer cells, compared to splenocytes from tumor naïve mice (fig 7E and F). The response levels between all tumor free mice were similar. The data supports that propranolol exerts antitumor effects that involve the formation of an anti-tumoral immune environment, and it suggests that combinations of propranolol and ICI could lead to durable therapeutic responses.

## Discussion

Retrospective analyses have shown a correlation between post-diagnostic use of non-selective beta blockers and increased relapse-free survival in breast cancer, melanoma, and prostate cancer (53,54). In this study we have demonstrated an anti-tumor effect of the non-selective beta blocker propranolol in a murine sarcoma model and a synergistic effect between propranolol and anti-CTLA4 checkpoint inhibitor therapy.

Propranolol is successfully used for treating high risk cases of proliferative infantile hemangioma, a benign vascular tumor, and has become the gold standard therapy due to its effectiveness and low toxicities (55). Propranolol’s suggested anti-angiogenic and anti-proliferative effects have been studied in multiple cancer types (18), but little is known about the effects of propranolol on STS. In this study, propranolol, as a single agent treatment, consistently resulted in a small but significant reduction in tumor growth and consequently an increased survival rate. One recent study indicates that human sarcoma cells express ADRB1, 2, and 3, and that their proliferation is inhibited by propranolol treatment *in vitro* (56). Similarly, we observed a dose dependent anti-proliferative effect of propranolol on MCA205 sarcoma cells *in vitro*. However, we did not see any propranolol-mediated inhibition of proliferation *in vivo* based on Ki67 immunostaining of tumor sections, indicating that the main mechanism of action was unlikely to be a direct effect on cancer cell proliferation. The absence of an anti-proliferative effect *in vivo* could be due to the intratumoral level of propranolol not reaching a sufficiently high concentration. The therapeutic potential of propranolol has also been attributed to an anti-angiogenic effect in breast cancer and vascular sarcomas (57,58). Here, we show that blocking of ADRB signaling by propranolol inhibits angiogenesis in MCA205 fibrosarcoma. This was reflected in a reduced gene expression of the pro-angiogenic molecule VEGFA as well as a striking reduction in vessel density in the tumors. The result suggests that the anti-angiogenic effect of propranolol can be extended to non-vascular STSs.

Studies on the therapeutic effect of propranolol have mainly focused on the direct effect on cancer cells, using immune deficient mice. However, the stimulation of ADRB2 on immune cells has been shown to dampen the inflammatory response towards infectious disease (59,60). In alignment with this effect, we observed changes in the immune cells of the TME upon propranolol treatment, in addition to the anti-angiogenic effect. Propranolol increased the number of tumor infiltrating CD4+ T cells and we also observed a trend of increased CD8+ T cells. We did not observe changes in the activation or exhaustion status of the T cells based on the expression of CD137 or PD1, respectively. Furthermore, we demonstrated that the full antitumor effect of propranolol was dependent on the presence of tumor-infiltrating T cells. T cell deficient nude mice, inoculated with MCA205 sarcoma cells, did not benefit from propranolol treatment in terms of tumor growth and survival rate.

In addition to the lymphoid compartment, we also observed changes in the myeloid compartment. Specifically, propranolol treatment resulted in a reduced number of infiltrating MDSCs in immune competent mice. The percentages of intratumoral MDSCs were similar between propranolol treated and control T cell deficient mice (supp. fig. S2C). This indicated that propranolol’s effect on the frequency of MDSCs was unlikely due to a direct modulation on MDSC recruitment and accumulation but rather a secondary effect that depends on the presence of T cells. In addition, we observed increased PD-L1 expression on TAMs in both immune competent and T cell deficient mice. Isolation of TAMs for RNA sequencing confirmed that propranolol induced a distinct gene expression profile, and an altered chemokine profile (supp. fig. S3D). Based on these changes, it remains a possibility that the changes in the lymphoid compartment could be partly caused by the altered TAM-mediated recruitment of lymphocytes. The direct impact of propranolol on macrophage function was confirmed *in vitro* using BMDMs. As observed *in vivo*, propranolol treatment increased the expression of PD-L1. ADRB stimulation by norepinephrine has been shown to induce M2-like polarization of macrophages (61). We confirmed the M2-polarizing effect of the ADRB agonist isoprenaline, while propranolol rescued the agonist induced downregulation of the M1-associated markers MHC-II and CD86. We do not exclude the possibility that propranolol could also modulate macrophage polarization through an ADRB independent effect. Propranolol has been shown to be a potent inhibitor of the phosphatidic acid phosphatase lipin-1 (62), which recently has been identified to be essential in IL4-mediated macrophage polarization *in vitro* (63).

In this study, we observed that propranolol treatment induces a shift of the TME towards a pro-inflammatory state, which is potentially more likely to respond to ICI therapies. A high number of tumor-infiltrating T cells has been associated with increased overall survival and response to immunotherapy in a variety of cancer types, including colorectal cancers, ovarian cancers, and melanoma (64,65). Moreover, the MDSC level has been identified as a promising biomarker for ICI treatment responses. A low number of MDSCs is associated with improved clinical response towards anti-CTLA4 and anti-PD1/PD-L1 checkpoint inhibitor therapies (66,67). In this study, we identified propranolol as a potent adjuvant to anti-CTLA4 treatment. By combining propranolol treatment with various ICI antibodies, we demonstrated that ADRB blockade significantly improves the efficacy of anti-CTLA4 therapy. The combination resulted in a reduced tumor growth rate and an increased overall survival. In one case, the combination treatment resulted in complete tumor regression. Subsequent re-challenge by inoculation of MCA205 cells and analysis of the presence of tumor-reactive splenocytes showed that the complete response was accompanied by acquired, long-lasting antitumor immunity. The acquired immunity was comparable to that achieved with combined anti-CTLA4 and anti-PD1 treatment. The combination of anti-CTLA4 and propranolol increased tumor infiltration of CD8+ T cells, compared to anti-CTLA4 alone. The anti-angiogenic function of propranolol was preserved, when used in combination with anti-CTLA4.

Few published studies have investigated the effect of ADRB blockade on the efficacy of ICIs. A retrospective analysis of melanoma patients treated with immune therapies showed that the concurrent usage of propranolol correlates with prolonged overall survival (68). Recently, a phase I clinical trial investigating the combination of propranolol and pembrolizumab in a limited number of patients indicated that the treatment combination is tolerable and could lead to increased response rates (69). Additionally, a preclinical study has shown that ADRB blockade by propranolol or genetic ablation of *Adrb2* improved anti-PD1 treatment outcome in a murine breast cancer model (70). In our study, propranolol treatment did not improve the response of MCA205 model to anti-PD1 or anti-PDL1 treatment. However, the MCA205 tumor model did generally respond very poorly to anti-PD1/PDL1 therapy, and it is possible that propranolol could enhance the efficacy of anti-PD1/PDL1 therapy in other tumor models, which are more responsive to this treatment. In the context of therapeutic cancer vaccines, propranolol has been demonstrated to improve naïve T cell priming, thereby boosting vaccine efficacy (71). In another study, the addition of anti-CTLA4 to a therapeutic cancer vaccine significantly improved the vaccine efficacy only when administered close to vaccination, highlighting the impact of anti-CTLA4 on priming of T cells in the lymphoid tissues (72). It is possible that propranolol and anti-CTLA4 synergistically stimulate the initial priming phase of an anti-cancer immune response, making this combination treatment particularly effective.

Immune therapies have given new possibilities of treatment to selected groups of cancer patients. However, many patients and cancer types remain unresponsive, and mechanisms limiting the treatment efficacy are yet to be fully elucidated (73). Combination treatment strategies that modulate the efficacy of immune therapies are thus attractive. Several preclinical studies have illustrated that psychological stress stimuli (e.g. social defeat, and social isolation) can promote tumor progression and metastasis through ADRB activation (74–76). Non-selective ADRB blockade, such as propranolol, has been proposed to reduce the impact of ADRB activation on disease outcome (18,77). In our study, the experimental mice were not subjected to any stress protocols, but housing temperature (∼22°C) of experimental mice, which is lower than the thermoneutral temperature for mice, has been identified as a source of adrenergic signaling (78). The chronic nature of this stimulus might resemble the stress faced daily by cancer patients.

The data presented in this study provide new insights into the inhibitory effect of propranolol on STS progression. In addition to an anti-angiogenic effect, propranolol modulates immune responses against STS by increasing effector cells and reducing the number of tumor-infiltrating immune suppressive cells such as MDSCs. The study supports the idea that adrenergic signaling contributes to therapy resistance and strengthens the rationale of combining beta blockers such as propranolol with ICI therapies for treating cancer patients.

## Author contributions

Conceptualization, N.J. and D.H.M.; Methodology, K.Y.F., L.G., N.J., and D.H.M.; Investigation, K.Y.F., A.M.A.R, V.G., A.Z.J., M-L.T., M.C., L.H.E., L.G., N.J., and D.H.M; Writing – Original Draft, K.Y.F., N.J., and D.H.M.; Writing – Review & Editing, K.Y.F., A.M.A.R, V.G., A.Z.J., M-L.T., M.C., L.H.E., L.G., N.J., and D.H.M.

