## Supplementary figures 1-4 and supplementary tables 1-2 for "Propranolol reduces sarcoma growth and enhances the response to anti-CTLA4 therapy by modulating the tumor microenvironment"

Supplementary figure S1

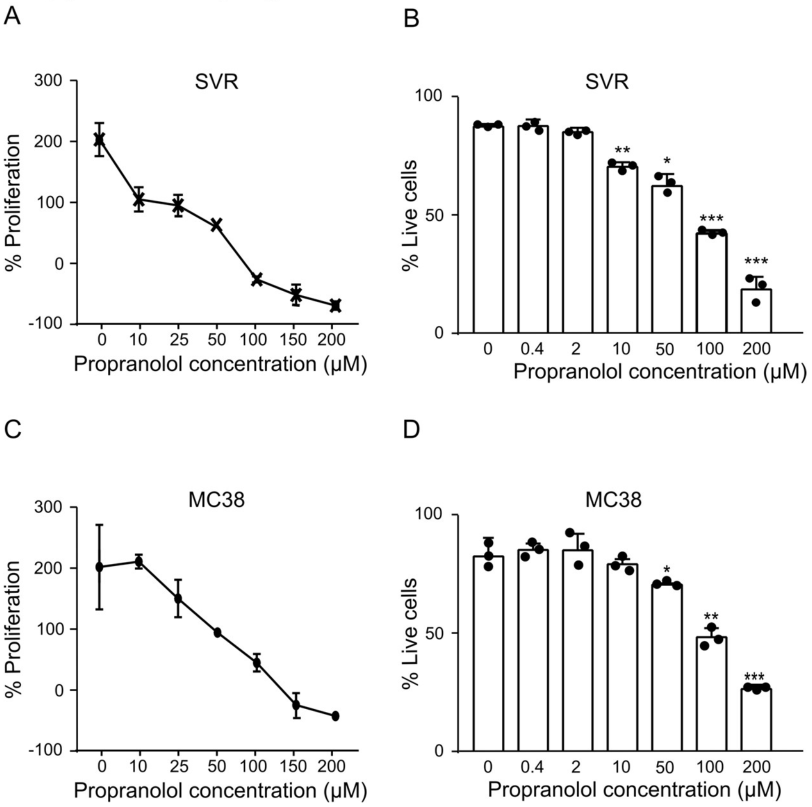


**Figure S1**. SVR sarcoma cells and MC38 carcinoma cells are sensitive to propranolol treatment. Dead cells were identified using trypan blue. Cell numbers are quantified manually using hemocytometer. **A-D,** Proliferation (**A and C**) and viability (**B and D**) of SVR **(A and B)** or MC38 cells **(C and D)** after 24 hours of propranolol stimulation at various concentrations (n=3). *p<0.05, **p<0.01, ***p<0.005, according to multiple t test with Bonferroni correction. Mean ± SD are depicted.

Supplementary figure S2

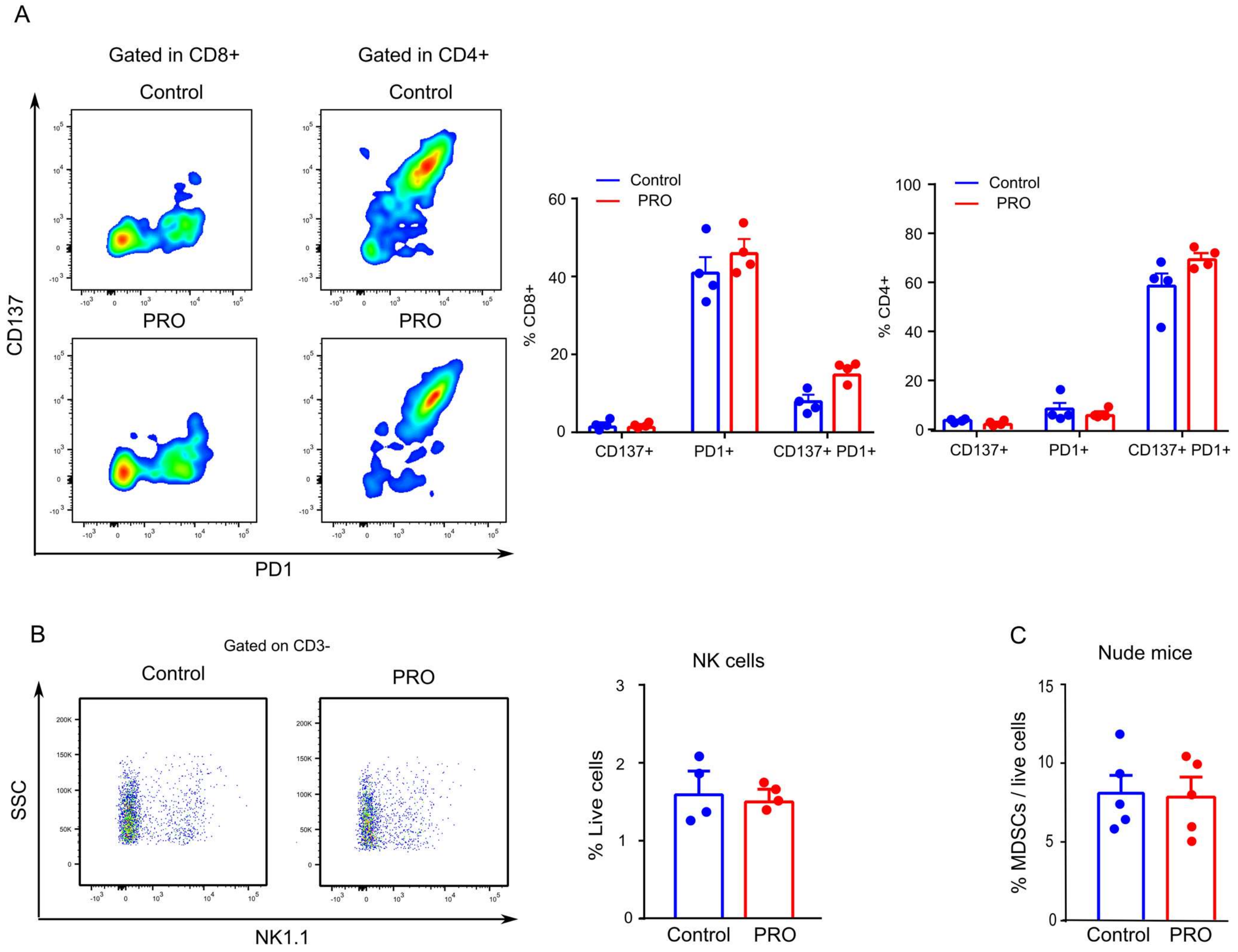


**Figure S2**. Effect of propranolol on the immune cells in the TME. **A-B,** Single cell suspensions were made from excised MCA205 tumors in C57BL/6 mice at the experimental endpoint and analyzed by flow cytometry (n=4). **A,** Representative flow cytometry dot plot (left) and quantification (right) of PD1 and CD137 expression on tumor infiltrating CD4+ or CD8+ T cells. **B,** Representative flow cytometry dot plot (left) and quantification (right) of NK cells in the TME. Single cell suspensions were made from excised MCA205 tumors in nude mice at endpoint and analyzed by flow cytometry (n=5). **C,** Quantification of intratumoral MDSCs (CD11b+ F4/80- GR1+) in MCA205 tumors grown in nude mice. Multiple t tests with Bonferroni correction for multiple comparison were used. Mean ± SEM are depicted.

Supplementary figure S3

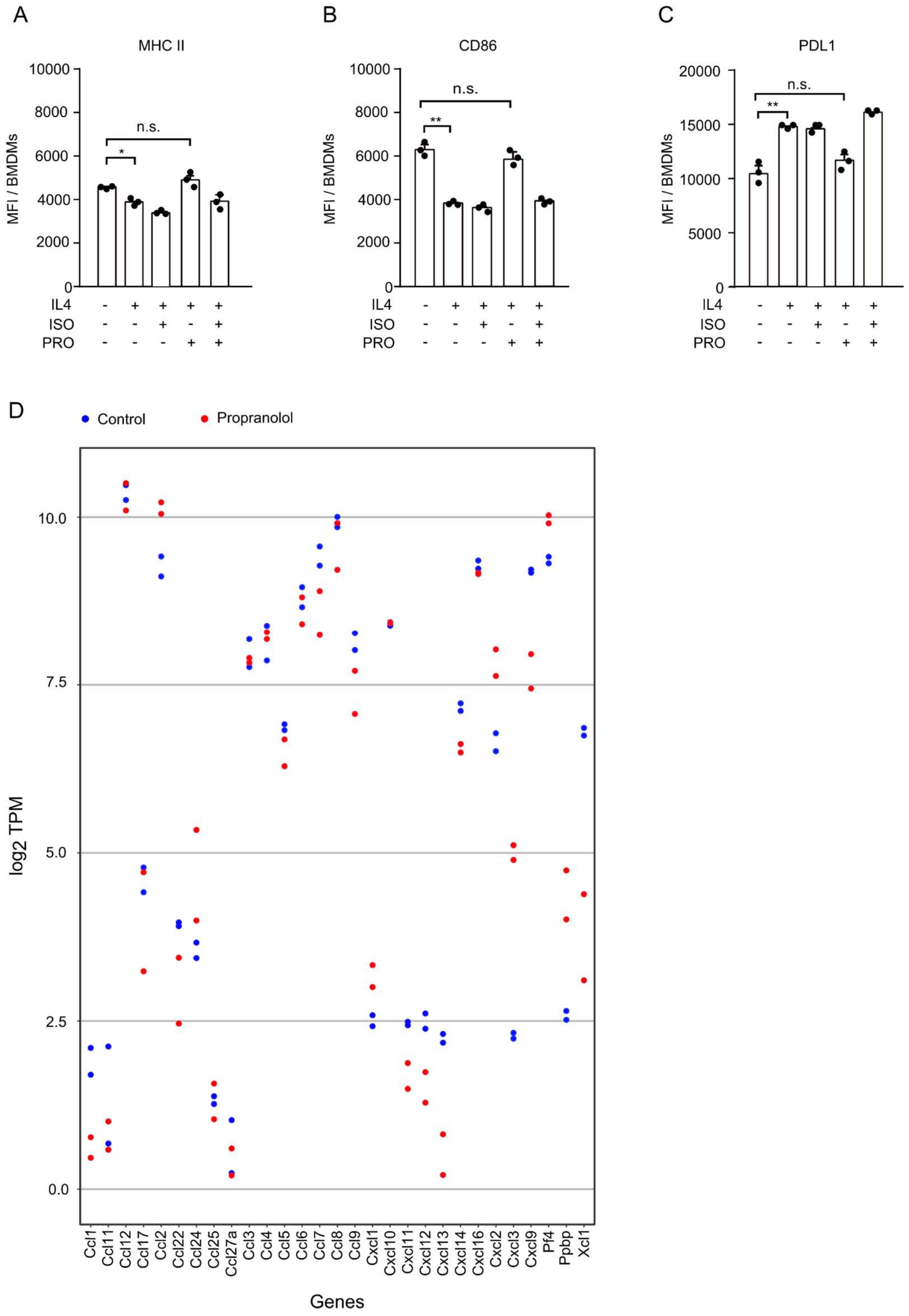


**Figure S3**. . Effect of propranolol on BMDMs and TAMs. **A-C,** BMDMs were stimulated with isoprenaline (ISO) or propranolol (PRO), in the presence of IL4, and cell surface marker expressions were evaluated by flow cytometry. Quantification of MHC II (**A**), CD86 (**B**), PDL1 (**C**) expression. **D,** TAMs from MCA205 tumors were sorted and subjected to RNA sequencing. **D,** The plot shows distinct chemokine expression in TAMs from the propranolol and control group. *p<0.05, **p<0.01, n.s. not significant, according to multiple t test with Bonferroni correction for multiple comparison. Mean ± SD are depicted.

Supplementary figure S4

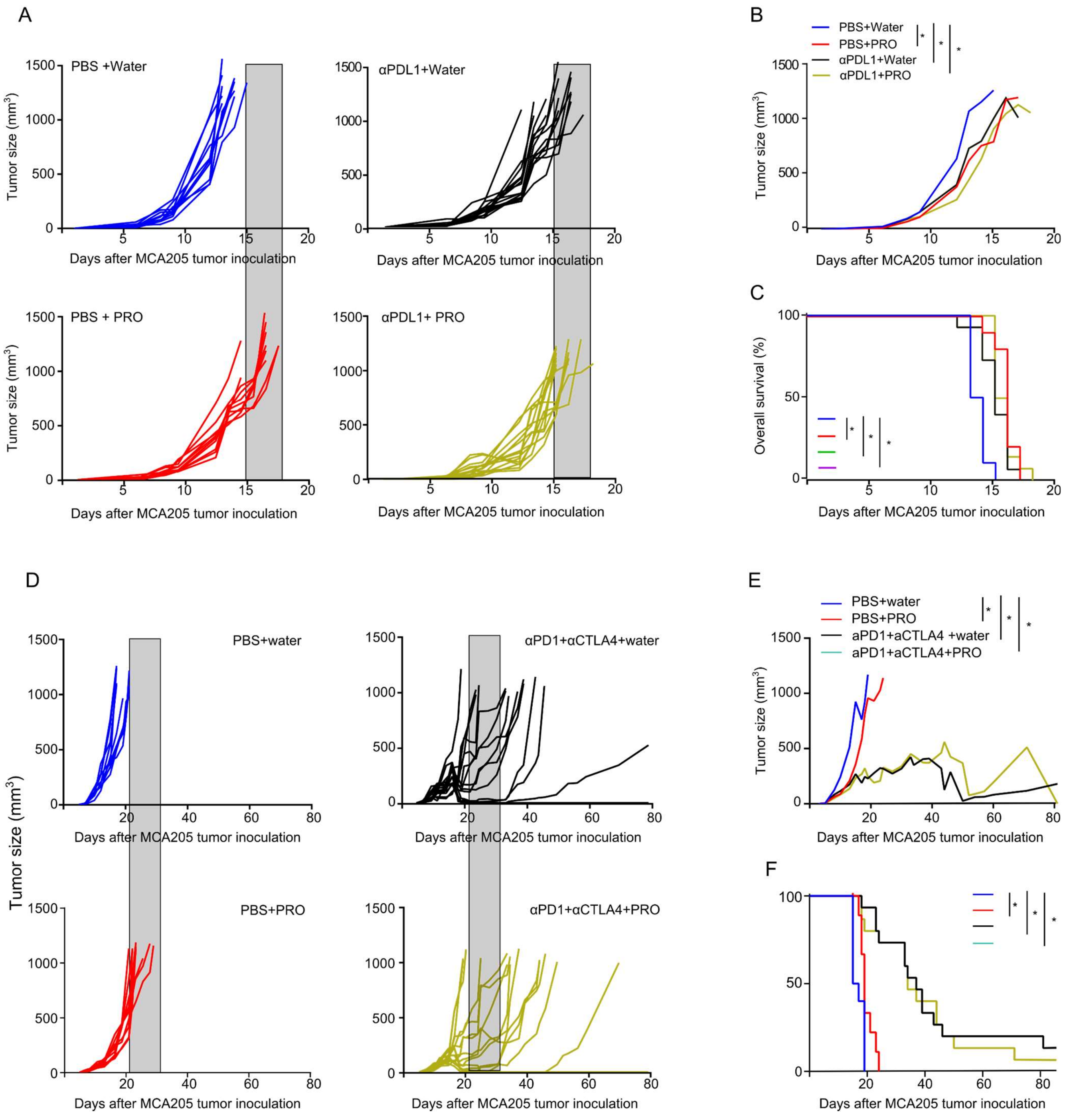


**Figure S4.** ADRB blockade by propranolol does not improve the treatment efficacy of anti-PDL1 and does not affect the efficacy of anti-PD1+anti-CTLA4 treatment in the MCA205 tumor model. **A-C,** Tumor growth kinetics (**A**), average tumor sizes per group (**B**), and Kaplan-Meier survival curves (**C**) of C57BL/6 mice inoculated with MCA205 cancer cells, treated with anti-PDL1 and propranolol (PRO). **D-F,** Tumor growth kinetics (**D**), average tumor sizes per group (**E**), and Kaplan-Meier survival curves (**F**) of C57BL/6 mice inoculated with MCA205 cancer cells, treated with anti-PD1, anti-CTLA4 and propranolol (PRO). Each line represents one animal (n=10-15 per group). Shaded area represents the terminal tumor growth time frame of PBS+water group. Statistical analyses were performed using TumGrowth software. *p<0.05

### Supplementary table 1. Antibodies used for flow cytometry analyses

| Antibody | Manufacturer | Catalog # |
|---|---|---|
| CD4 BV421 | BioLegend | 100438 |
| CD8 APC | BioLegend | 100712 |
| CD3 FITC | BioLegend | 100204 |
| CD11b PE-Cy7 | BioLegend | 101215 |
| F4/80 APC | BioLegend | 123116 |
| CD206 PE | BioLegend | 141705 |
| PDL1 BV421 | BioLegend | 124315 |
| PD1 PE | BioLegend | 109103 |
| CD137 | eBioscience | 25-1371-82 |
| GR1 PerCP-Cy5.5 | BioLegend | 108427 |
| CD25 PerCP-Cy5.5 | BioLegend | 101912 |
| NK 1.1 APC-Cy7 | BioLegend | 108724 |
| CD86 PerCP-Cy5.5 | BioLegend | 105027 |
| MHC II APC-Cy7 | BioLegend | 107627 |
| CD45 FITC | eBioscience | 11-0451-82 |
| CD44 APC-Cy7 | BioLegend | 103027 |
| CD62L BV650 | BD Biosciences | 564108 |
| IFNy PE | BioLegend | 505808 |
| TNFa BV605 | BioLegend | 506329 |

Supplementary table 2. Primer sequences for qRT-PCR on MCA205 whole tumor RNA or MCA205 cell line

| **Gene name** | **5’ Forward 3** | **5’ Reverse 3’** |
|---|---|---|
| *Vegfa* | GGCCTCCGAAACCATGAACT | CTGGGACCACTTGGCATGG |
| *Kdr* | TTGTGAATGTCCCACCCCAG | TTGGCGTAGACTGTGCATGT |
| *Adrb1* | TCATCGTGGTGGGTAACGTG | ACCAGCAATCCCATGACCAG |
| *Adrb2* | TGGTTGGGCTACGTCAACTC | TCCGTTCTGCCGTTGCTATT |
| *Actb* | CACTGTCGAGTCGCGTCC | TCATCCATGGCGAACTGGTG |
